# FLASH radiotherapy drives intestinal regeneration via macrophage-derived WNT4 signaling and attenuated T-cell inflammation

**DOI:** 10.64898/2026.11.08.764105

**Authors:** Yixun Zhu, Yidan Wang, Yan Zhang, Chaofan Lin, Yucheng Wang, Chenyang Huang, Hao Zha, Tuchen Huang, Jian Wang, Caihui Li, Rui Qiu, Zhihui Li, Guofu Dong, Wei Wang, Hao Lu, Changzhen Wang, Qibin Fu

## Abstract

Ultra-high dose rate radiotherapy (FLASH-RT) reduces normal-tissue toxicity, yet how elevated dose rate improves tissue repair remains unclear. Here we combined whole-abdominal FLASH-RT and conventional radiotherapy (CONV-RT) in mice with single-cell transcriptomics, trajectory analysis, cell–cell communication inference, and in vivo perturbation experiments to define the intestinal repair ecosystem. Compared with CONV-RT, FLASH preserved Lgr5^+^ intestinal stem cells and Fabp1^+^ absorptive epithelial cells. Trajectory analysis showed that FLASH maintained stem-cell differentiation toward mature epithelial states enriched for fatty-acid metabolism. During a day-3 repair window, FLASH was accompanied by enrichment of Mrc1⁺ macrophages, increased WNT4-associated signaling involving Fzd5⁺Lrp5/6-expressing stem cells, and early transient β-catenin activation. In vivo augmentation of Wnt4 signaling promoted crypt regeneration after CONV-RT. FLASH also reduced Tcf7⁺ and Tbx21⁺ CD4⁺ T-cell populations and TNF-α levels, consistent with attenuation of the pro-inflammatory CD4⁺ T-cell response during intestinal recovery. Together, these findings indicate that FLASH-associated tissue sparing involves coordinated remodeling of regenerative and inflammatory immune states, suggesting that radiation dose rate influences not only initial tissue injury but also the immune microenvironment of subsequent recovery. Targeting this immune–stem cell crosstalk may provide a strategy to mimic FLASH-like radioprotection.

## Introduction

Ionizing radiation is a cornerstone of treatment for abdominal and pelvic malignancies. However, the intestine is highly radiosensitive because epithelial homeostasis depends on the continuous turnover of rapidly renewing stem and progenitor cells within the crypts ^1–3^. Radiation depletes these crypt cells, disrupts epithelial integrity, and triggers inflammation, culminating in crypt loss, villus atrophy, mucosal injury, and, in severe cases, gastrointestinal syndrome ^4–6^. These toxicities limit dose escalation, making normal-tissue preservation a central challenge in radiotherapy. Regeneration after injury depends on both the survival of resident stem cells and the local microenvironment that guides their response to damage ^7–9^. In the intestine, Lgr5⁺ crypt base columnar (CBC) cells fuel epithelial turnover during homeostasis, whereas reserve and injury-responsive cell populations drive epithelial regeneration after severe damage ^10–12^. The intestinal stem cell niche integrates epithelial, mesenchymal, vascular, and immune components that collectively regulate stem cell maintenance, proliferation, and differentiation ^13–16^. Irradiation dynamically remodels this niche, with inflammatory and regenerative programs unfolding in parallel. Thus, tissue recovery is shaped not merely by which stem cells survive, but by the cellular environment that supports their subsequent regeneration.

Ultra-high dose rate radiotherapy (FLASH-RT) (>40 Gy/s) markedly reduces normal-tissue toxicity while maintaining tumor control, a phenomenon commonly referred to as the “FLASH effect” ^17–21^. Several studies have implicated alterations in redox responses and antioxidant defenses in FLASH-mediated tissue protection ^22^. More recent studies have focused on intestinal stem cell populations. FLASH-RT has been reported to preserve Lgr5⁺ CBCs and Bmi1-associated quiescent or reserve stem cell populations. Moreover, single-cell RNA sequencing (scRNA-seq) of proton FLASH-treated intestines identified preserved Lgr5⁺ CBCs and injury-responsive regenerative stem cells (revSCs) characterized by markers including Ly6a, Clu, Areg, and Anxa2 ^23^. These studies also revealed changes in cell-cycle states within these populations, suggesting that FLASH-RT can influence the cellular state of surviving regenerative cells. However, whether FLASH-associated stem-cell preservation reflects coordinated remodeling of the multicellular regenerative niche remains poorly understood..

Immune cells are integral to tissue recovery after radiation injury, coordinating inflammatory responses with epithelial regeneration through signals that regulate stem and progenitor cells ^24–28^. Studies of ionizing radiation have shown that radiation dynamically remodels the intestinal immune microenvironment, with distinct immune populations undergoing differential depletion, recruitment, and recovery ^29,30^. Single-cell studies further revealed shifts in monocyte/macrophage states and T-cell subsets, together with altered immune-cell communication during intestinal injury and regeneration ^30,31^. FLASH preserves intestinal crypts and stem cells while attenuating radiation-induced inflammation, with recent single-cell studies also revealing changes in macrophage and T-cell abundance ^20,32^. However, these studies have largely characterized broad inflammatory responses or changes in immune-cell composition, leaving the functional organization of specific immune states and their communication with regenerating epithelium poorly defined. Importantly, radiation dose rate may influence not only initial tissue damage, but also which cellular populations persist, which regenerative signals emerge, and how immune cells interact with surviving stem cells. Defining these dose-rate-dependent changes may reveal how FLASH shapes the regenerative trajectory beyond initial radiation injury.

Here, we combined single-cell transcriptomic profiling, multiplex immunohistochemistry, and functional validation to define the cellular and molecular basis of intestinal responses to FLASH-RT and conventional radiotherapy (CONV-RT). We found that FLASH-RT induced coordinated myeloid and lymphoid changes during tissue recovery. In the myeloid compartment, FLASH-RT enriched Mrc1⁺ macrophages and was associated with WNT4 communication with Olfm4⁺ intestinal stem cells, accompanied by transient activation of β-catenin signaling during the early regenerative phase. In parallel, FLASH-RT was associated with reduced Tbx21⁺ CD4⁺ T-cell abundance and diminished TNF-α production, consistent with a less inflammatory environment during epithelial recovery. These findings support a model in which FLASH-RT sparing effect not only preservation of regenerative epithelial populations but also remodeling of the immune–stem cell niche that shapes recovery. More broadly, our findings suggest that radiation dose rate can be an important regulator of the multicellular environment in which repair unfolds.

## Results

### Dynamic Single-Cell Landscape of the Small Intestine Following CONV-RT and FLASH-RT

To systematically characterize the effects of FLASH-RT on intestinal injury and repair, we established murine whole-abdominal irradiation models using CONV-RT and FLASH-RT at 15 Gy (Fig. 1A) ^22^, a dose at which FLASH-RT substantially attenuates intestinal injury. Mice were analyzed at 3 and 6 days after irradiation, corresponding to the acute injury phase and the subsequent crypt regeneration and epithelial reconstruction phase, respectively ^33–35^.

**Fig. 1.**
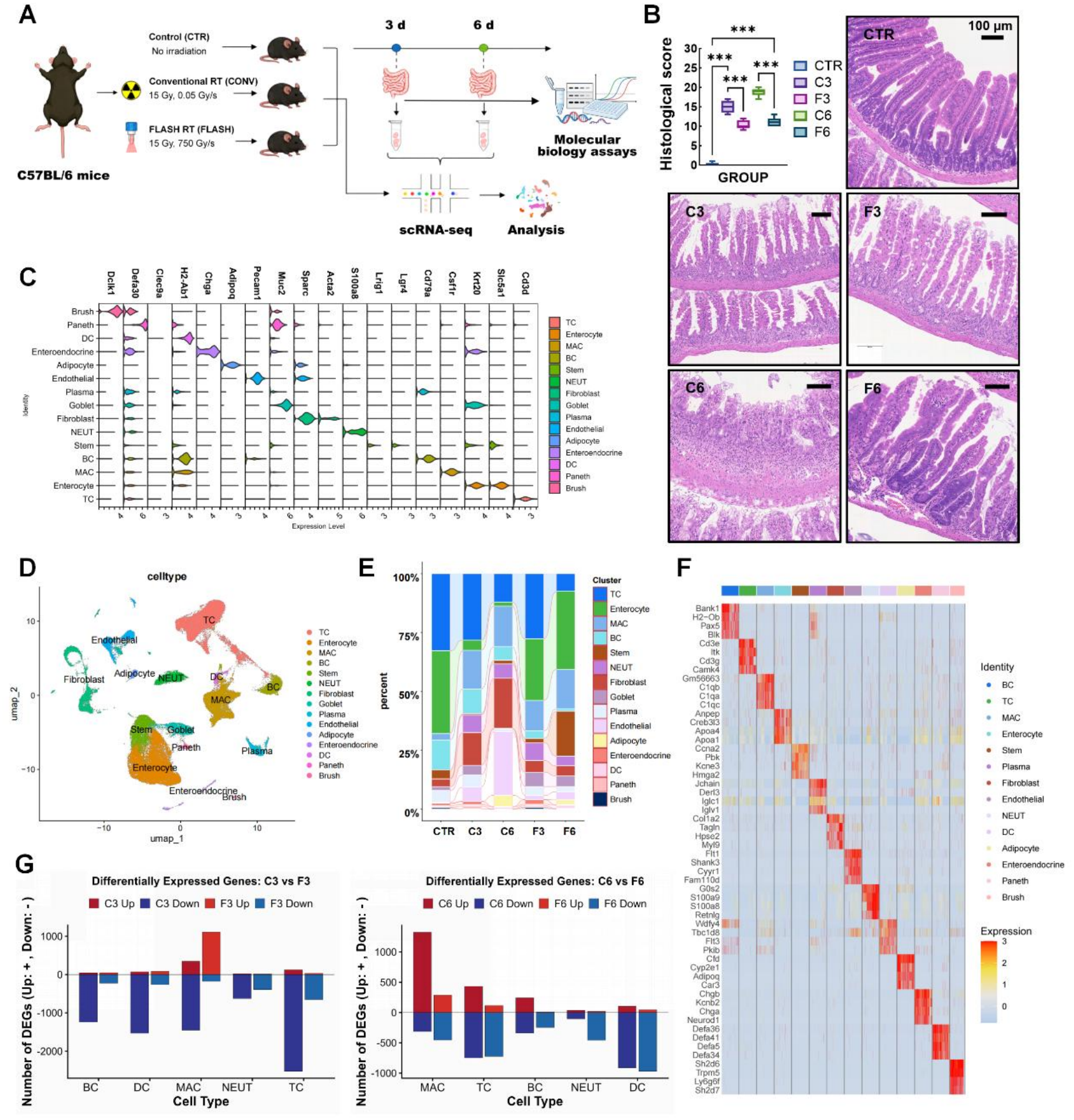
FLASH-RT reshapes the cellular landscape of the regenerating intestine after radiation injury. (A) Schematic of the experimental design and sampling strategy. (B) Representative H&E-stained sections of the small intestine from CTR, C3, F3, C6, and F6 groups (n = 9 from three mice per group). (C) Violin plots showing representative marker gene expression for the annotated cell populations. (D) UMAP visualization of the major intestinal cell populations. (E) Relative abundance of major intestinal cell populations across experimental groups shown as a river plot. (F) Heatmap of representative marker genes across the identified cell populations. (G) Bar plot denoting the number of DEGs in each group on Day 3 and Day 6 compared to CTR. Data are presented as mean ± SEM. Statistical significance was assessed by a one-way ANOVA followed by Tukey’s multiple comparisons test. *\*P* < 0.05, *\*\*P* < 0.01, *\*\*\*P* < 0.001. Scale bars are indicated.

Histological and proliferation analyses revealed modality-specific tissue recovery. At day 3 after CONV-RT (C3), H&E staining (Fig. 1B) showed marked crypts loss,substantial disruption of the remaining crypt architecture, pronounced villus shortening, and epithelial denudation. Consistent with these histological changes, Ki67 immunofluorescence showed a marked reduction in proliferative activity in C3 compared with CTR, whereas F3 exhibited significantly higher Ki67-positive area than C3. By day 6 (C6), partial crypt regeneration and villus restoration were evident, but overall tissue recovery remained limited. In contrast, FLASH-RT preserved more intact crypts at day 3 (F3), with better epithelial continuity than C3. At day 6, Ki67-positive area was significantly higher in F6 than in C6, although it remained below the baseline level in CTR. Consistent with these findings, crypt numbers and villus length were significantly greater in F6 than in C6 and approached baseline. These findings confirm FLASH-RT attenuates acute intestinal injury and enhances subsequent crypt regeneration relative to CONV-RT.

We then performed scRNA-seq on 14 samples across five groups (CTR, C3, F3, C6, F6). After quality control and Harmony batch correction, 120,633 high-quality cells (mean 1,577 genes and 5,059 UMIs per cell; 25,985 genes detected) underwent unsupervised clustering. Based on the expression of canonical marker genes, we annotated 15 distinct cell types (Fig. 1C–F), comprising five intestinal epithelial cell (IEC) populations, one progenitor-cell population, three lymphoid populations, three myeloid populations, and three mesenchymal/stromal cell populations.

Cell-type composition analysis revealed extensive remodeling of the intestinal cellular landscape after both CONV and FLASH-RT (Fig. 1E). Among intestinal epithelial cells, enterocytes represented the predominant population and were more abundant after FLASH-RT than CONV-RT. A similar pattern was observed for the stem cell population within the progenitor compartment. In contrast, the overall proportions of macrophages and T cells were broadly comparable between FLASH-RT and CONV-RT at the major-cell-population level. Nevertheless, both populations exhibited substantial transcriptional differences between irradiation modalities and across time points, as reflected by the number and composition of differentially expressed genes. (Fig. 1G). These observations prompted us to examine whether FLASH-RT differentially affects epithelial regenerative states and immune-cell states during intestinal recovery.

### FLASH-RT Preserves Olfm4⁺ Stem Cells and Promotes Enterocytic Recovery during Intestinal Regeneration

To dissect the cellular dynamics of intestinal regeneration following FLASH-RT, subpopulation analysis was performed, identifying four stem cell subsets with distinct transcriptional signatures and four epithelial cell subsets at different differentiation stages (Fig. 2A, 2B). Given the marked enrichment of the Olfm4⁺ stem cell subset (Stem1) and Epcam⁺ Fabp1⁺ mature enterocytes (Enterocyte1) after FLASH-RT (Fig. 2C), we focused on these two cell populations and combined multiplex immunohistochemistry (mIHC), functional enrichment analysis and pseudo-time trajectory inference to dissect FLASH-specific regeneration.

**Fig. 2.**
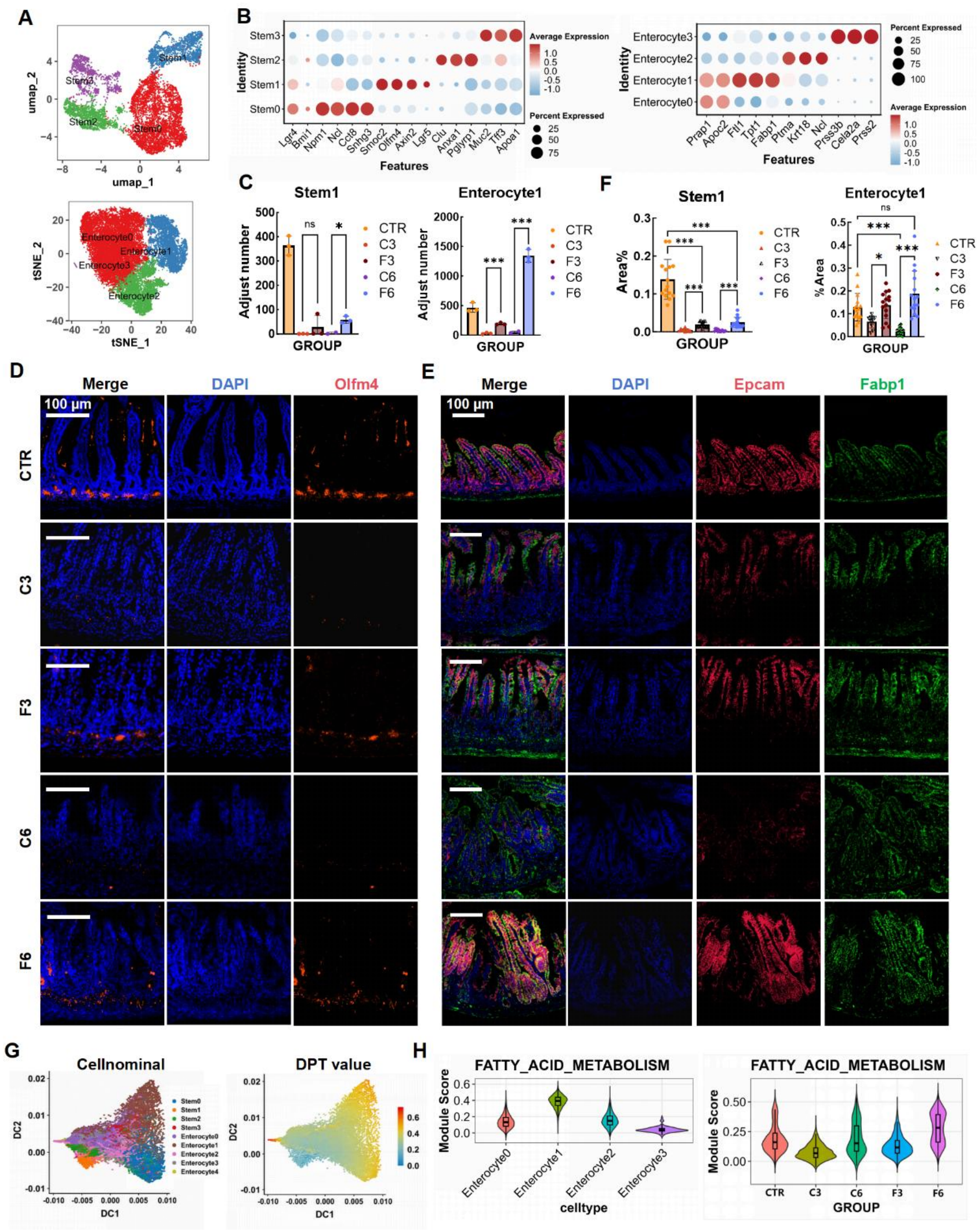
FLASH-RT preserves intestinal stem-cell populations and promotes regenerative epithelial states. (A) UMAP visualization of intestinal stem-cell and enterocyte populations. (B) Dot plots showing representative marker gene expression in intestinal stem-cell and enterocyte populations. (C) Quantification of intestinal stem-cell and enterocyte populations across the indicated experimental groups (n = 3). (D) Representative multiplex immunofluorescence images showing intestinal stem-cell populations in the indicated experimental groups. Scale bar = 100 μm. (E) Representative multiplex immunofluorescence images showing enterocyte populations in the indicated experimental groups. Scale bar = 100 μm. (F) Quantification of intestinal stem-cell populations by multiplex immunofluorescence (n = 15 from three mice per group). (G) Pseudotime trajectory analysis of intestinal epithelial cells. DPT, diffusion pseudotime. (H) Violin plots of Gene set analysis of intestinal stem-cell and enterocyte populations. Data are presented as mean ± SEM. Statistical significance was assessed by a two-sided unpaired Student’s *t* test or one-way ANOVA followed by Tukey’s multiple comparisons test. *\*P* < 0.05, *\*\*P* < 0.01, *\*\*\*P* < 0.001. Scale bars are indicated.

Quantitative mIHC results (Fig. 2D, 2F) showed that Stem1 cells predominated at the crypt base in control group (CTR). At day 3 post-irradiation, both C3 and F3 groups exhibited a substantial decrease in stem cell proportion relative to CTR, but F3 group retained significantly more than C3 group, suggesting that FLASH confers a protective advantage to stem cells as early as the acute phase. By day 6, this divergence was further widened. C6 group showed further decline, whereas F6 exhibited marked Stem1 enrichment. These findings indicate greater preservation of the Olfm4⁺ stem-cell compartment during the acute phase and more robust stem-cell recovery following FLASH-RT. Enterocyte1 quantification (Fig. 2E, 2F) revealed that F6 group attained epithelial area fractions comparable to CTR (ns) and significantly exceeding C6, whereas F3 and C3 did not differ (ns); the Enterocyte1 proportions in the C3 and C6 groups remained consistently low throughout. These findings indicate enhanced recovery of the mature enterocyte compartment following FLASH-RT, with the most pronounced difference emerging by day 6.

To examine the relationship between stem-cell recovery and enterocytic differentiation, we performed pseudotime analysis. Stem1 occupied an early position along the inferred differentiation trajectory, which progressed toward Enterocyte1 and was accompanied by concordant changes in lineage-associated marker genes (Fig. 2G). We next characterized the transcriptional features associated with mature epithelial recovery. GO enrichment analysis and gene-set scoring showed enrichment of fatty acid metabolism-related programs in mature epithelial populations after FLASH-RT, with the highest activity at day 6 (Fig. 2H). Together with their increased relative abundance in F6, these transcriptional features were consistent with recovery of a differentiated enterocyte state after FLASH-RT.

Collectively, these findings delineate a temporally coordinated epithelial regenerative response following FLASH-RT, characterized by greater preservation of Olfm4⁺ stem cells during the acute injury phase, robust recovery of the stem-cell compartment by day 6, and subsequent restoration of mature enterocyte populations with metabolic features characteristic of differentiated intestinal epithelium. The inferred stem-to-enterocyte trajectory further supports a link between stem-cell recovery and epithelial reconstruction after FLASH-RT.

### FLASH-RT Enriches Mrc1⁺ Macrophage with Anti-Inflammatory and Pro-Repair Phenotypes

To investigate the dynamic remodeling of macrophages following FLASH-RT and their potential involvement in intestinal repair, we performed macrophage subpopulation analysis and identified five transcriptionally distinct subsets (Fig. 3A, 3B). Comparison of macrophage subset composition revealed a marked enrichment of MAC2 in the FLASH-RT groups relative to the corresponding CONV-RT groups. (Fig. 3C). This subset was characterized by high Mrc1 expression and was therefore referred to as Mrc1⁺ MAC. To characterize the transcriptional phenotypes of these macrophage subsets, we calculated M1 and M2 gene-set scores. Mrc1⁺ MAC showed the highest anti-inflammatory score, whereas Ly6c2⁺ MAC0 showed the highest pro-inflammatory score (Fig. 3D). Gene Ontology (GO) analysis (Fig. 3E) further associated Mrc1⁺ MAC with tissue repair, apoptotic cell phagocytosis, and inflammation resolution pathway.

**Fig. 3.**
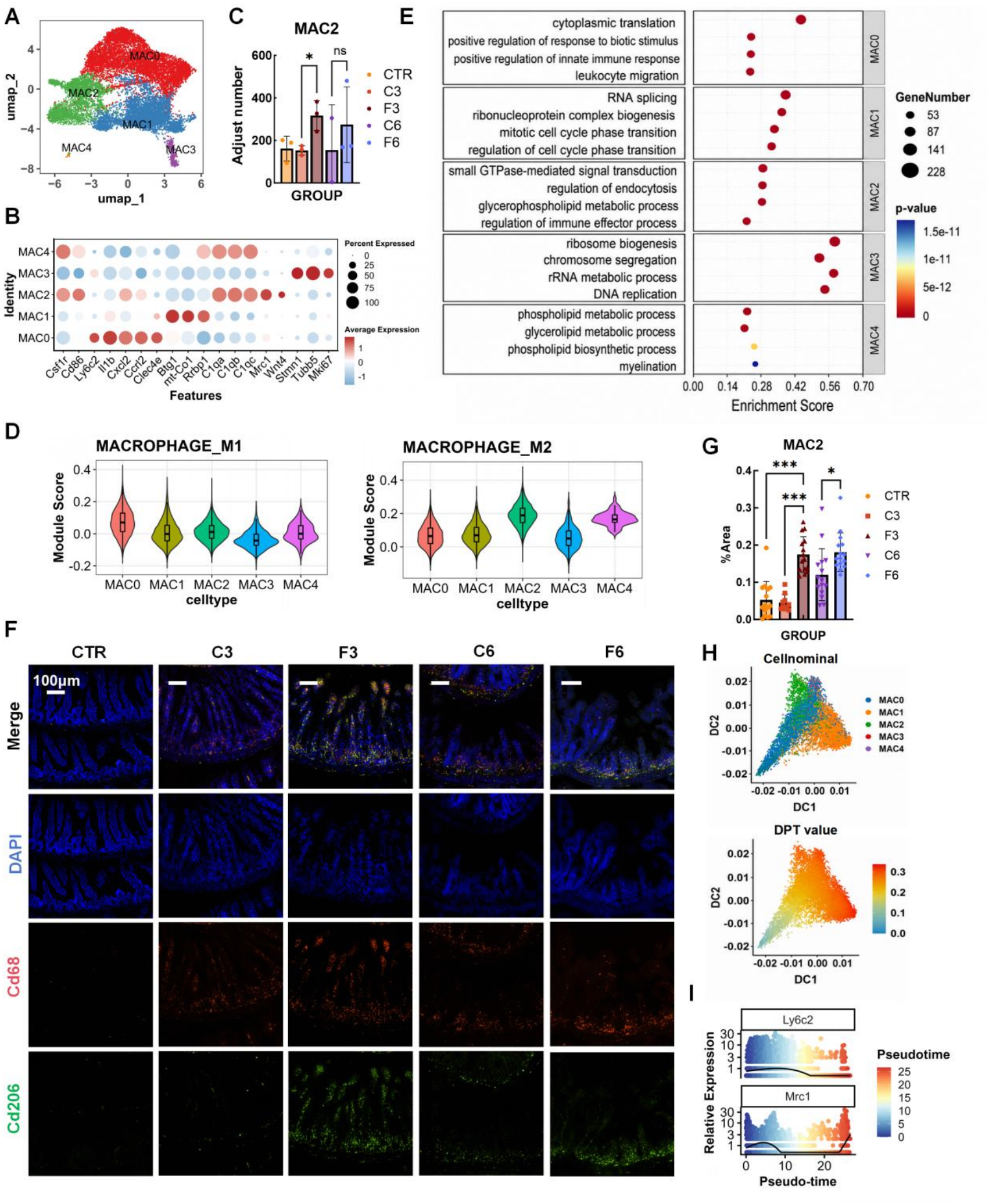
FLASH-RT remodels macrophage states during intestinal regeneration. (A) UMAP visualization of intestinal macrophages identified by scRNA-seq. (B) Dot plot showing representative marker gene expression across the identified macrophage populations. (C) Quantification of macrophage populations across the indicated experimental groups (n = 3). (D) Gene set analysis of macrophage subpopulations, shown as violin plots. (E) Gene Ontology (GO) enrichment analysis of differentially expressed genes in macrophage populations. (F) Representative multiplex immunofluorescence images showing macrophage populations in the indicated experimental groups. Scale bar = 100 μm. (G) Quantification of macrophage populations by multiplex immunofluorescence (n = 15 from three mice per group). (H) Pseudotime trajectory analysis of macrophage populations. DPT, diffusion pseudotime. (I) Expression dynamics of Ly6c2 and Mrc1 along the macrophage pseudotime trajectory. Data are presented as mean ± SEM. Statistical significance was assessed by a one-way ANOVA followed by Tukey’s multiple comparisons test. *\*P* < 0.05, *\*\*P* < 0.01, *\*\*\*P* < 0.001. Scale bars are indicated.

To validate these findings *in situ*, we performed precise quantification of the spatial distribution and abundance of Mrc1⁺ macrophages in the intestinal mucosa using multiplex immunohistochemistry (mIHC). Results (Fig. 3F, 3G) showed low and comparable area fractions in CTR and C3 group, but marked elevation in F3 and F6 group relative to matched CONV time points. These findings confirmed the preferential accumulation of Mrc1⁺ macrophages in FLASH-irradiated intestines during the regenerative period.

To further elucidate the potential state transitions and polarization dynamics of Mrc1⁺ macrophages, we constructed pseudotime differentiation trajectories of macrophage subsets using Monocle and diffusion map-based approaches. This analysis revealed a continuous spectrum from pro-inflammatory phenotype toward anti-inflammatory/pro-repair phenotype (Fig. 3H). MAC0 (high proinflammatory score) was specified as the trajectory root, whereas Mrc1⁺ MAC (high anti-inflammatory score) localized to the terminal end. Along pseudotime (Fig. 3I), pro-inflammatory signature gene Ly6c2 declined while anti-inflammatory/pro-repair signature gene Mrc1 rose, consistent with a transcriptional transition from an inflammatory state toward an anti-inflammatory/pro-repair state.

Collectively, these findings demonstrate that FLASH-RT enriches Mrc1⁺ anti-inflammatory macrophages by day 3 and sustains them through day 6, establishing a pro-repair, inflammation-resolving microenvironment. This immune remodeling coincides temporally with Olfm4⁺ stem cells regeneration and epithelial differentiation, consistent with a model in which Mrc1^+^ macrophages provide niche signals that support stem cell survival, expansion, and differentiation.

### Mrc1⁺ Macrophages Engage WNT4–β-Catenin Signaling during FLASH-Associated Intestinal Regeneration

To identify the signaling mediators between macrophages and stem cells, we performed cell–cell communication analysis on the scRNA-seq data. We found that overall communication activity of the WNT pathway was higher in F3 than C3 group (Fig. 4A). Cell–cell roles dissection (Fig. 4B) identified MAC2 macrophages as the primary WNT senders and Stem1 as the main receivers, indicating directional WNT transmission from macrophages to stem cells.

**Fig. 4.**
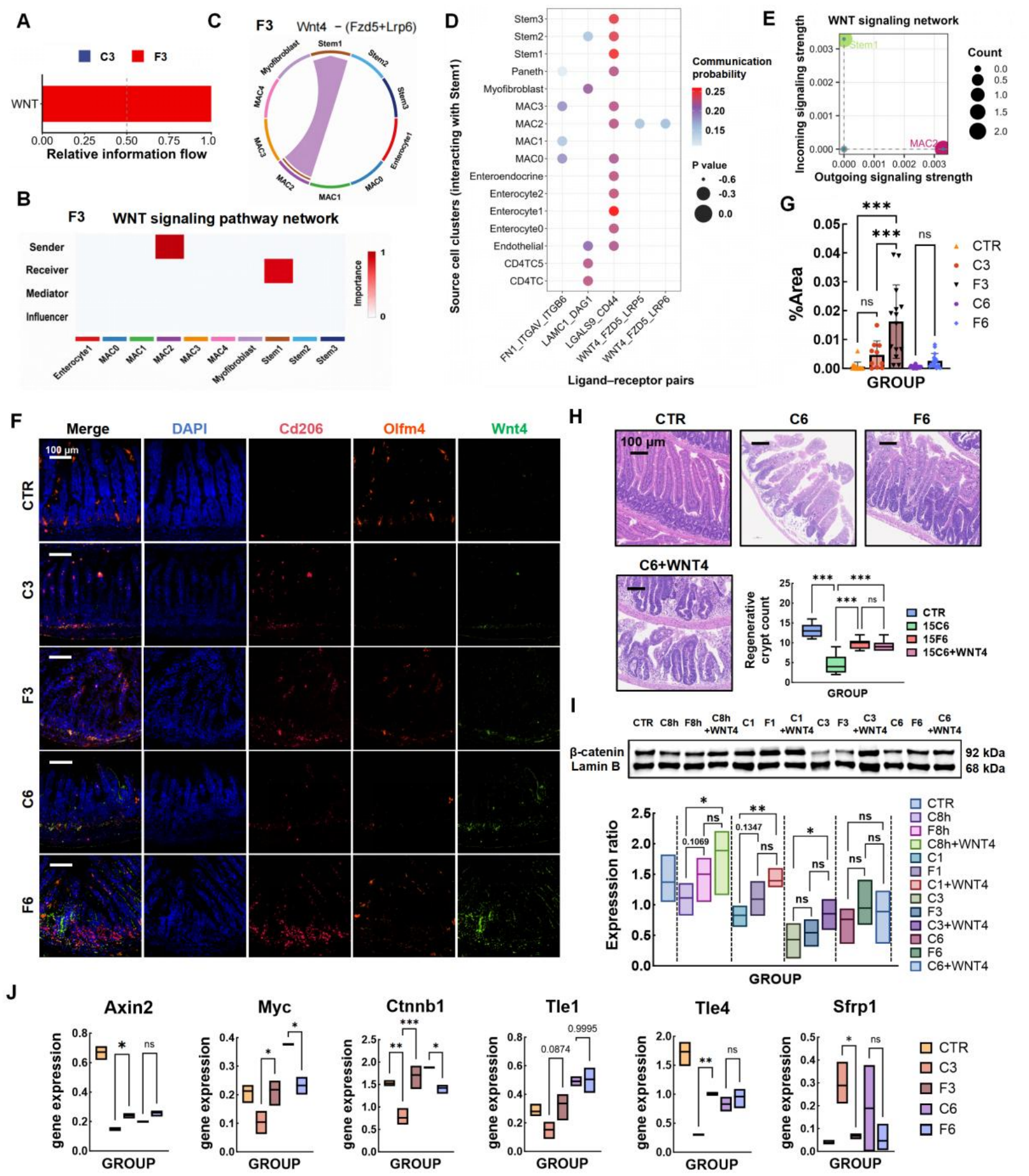
FLASH-RT enhances macrophage–stem-cell WNT4 signaling and promotes intestinal regeneration. (A) Relative information flow of WNT signaling between the C3 and F3 groups identified by CellChat analysis. (B) WNT signaling network in the F3 group, showing the major signaling sources and target cell populations. (C) Cell–cell communication network showing WNT4–FZD5/LRP6 signaling between macrophages and target cells in the F3 group. (D) Differential ligand–receptor interactions between the C3 and F3 groups involving Stem1 cells. (E) Major signaling inputs and outputs within the WNT signaling network. (F) Representative multiplex immunofluorescence images showing macrophage populations in the indicated experimental groups. (G) Quantification of macrophage populations by multiplex immunofluorescence (n = 15 from three mice per group). (H) Representative H&E-stained sections of the small intestine from CTR, C6, F6, and C6 + WNT4 groups. And quantification of intestinal regenerative features in the indicated groups. (I) Western blot analysis of nuclear active β-catenin in intestinal tissues. Lamin B was used as the loading control (n=4). (J) Single-cell RNA-seq analysis of β-catenin downstream target gene expression (n=3). Data are presented as mean ± SEM. Statistical significance was assessed by a unpaired Student’s *t* test or one-way ANOVA followed by Tukey’s multiple comparisons test. *\*P* < 0.05, *\*\*P* < 0.01, *\*\*\*P* < 0.001. Scale bars are indicated.

To elucidate the underlying molecular interactions, we focused on the binding of Wnt4 ligand to the Fzd5–Lrp receptor complex. Chord diagram (Fig. 4C) showed strong Wnt4–(Fzd5+Lrp6) and Wnt4–(Fzd5+Lrp5) signaling from MAC2 to Stem1 in the F3 group. Quantitative analysis of signaling probabilities for each cell–cell pair (Fig. 4D) further confirmed that Wnt4 signal transmission from MAC2 to Stem1 exhibited the highest interaction strength exclusively in the F3 group. Outgoing/incoming signaling scatter plots (Fig. 4E) highlighted distinct signaling patterns, with Stem1 exhibiting high incoming and relatively low outgoing signaling, whereas MAC2 showed the opposite pattern. Together, these data indicate that FLASH-RT enhances the macrophage–stem cell WNT communication axis via Wnt4–Fzd5/Lrp.

To visually validate the WNT signaling communication predicted by single-cell analyses at the tissue level, we performed mIHC co-staining for the MAC2 marker Mrc1, the intestinal stem cell marker Olfm4, and the WNT ligand WNT4 (Fig. 4F), and quantified the colocalization area fraction (%Area) among these three markers. The colocalization area of Cd206/Olfm4/WNT4 in the F3 group was markedly higher than that in CTR and C3 group (Fig. 4G), while remaining low in the CTR, C3, C6, and F6 groups. These histological data demonstrate spatial enrichment of WNT4 at Mrc1+ macrophage–Olfm4+ stem cell interfaces in F3, supporting paracrine WNT4 signaling. Western blot analysis further showed that the WNT4 protein level was higher in bone marrow-derived macrophages from FLASH-irradiated mice than those from CONV-irradiated groups.

To test WNT4 function in vivo, we used AAV-mediated WNT4 overexpression (C6+WNT4). Quantitative HE staining (Fig. 4H) showed that crypt numbers were reduced in C6 relative to CTR (P < 0.0001) but restored in both F6 and C6+WNT4 (ns vs each other; both higher than C6), indicating that WNT4 overexpression is sufficient to promote crypt regeneration after CONV irradiation to an extent comparable to FLASH.. Given that nuclear β-catenin was unchanged at day 6 (Fig. 4I), we examined earlier time points by Western blot: β-catenin was elevated in FLASH and CONV+WNT4 relative to CONV at 8 h, 1 d, and 3 d, returning to baseline by day 6, an early-activation, late-decline pattern consistent with prior work ^36^. scRNA-seq data (Fig. 4J) further confirmed that WNT/β-catenin target genes (Axin2, Ctnnb1, Myc, Tle1, Tle4) were up-regulated in Stem1 at day 3 in FLASH and CONV+WNT4 but declined by day 6, whereas the antagonist Sfrp1 showed the opposite trend.

Together, these data show that WNT4 is sufficient to promote crypt regeneration after CONV-RT and that FLASH-RT is associated with WNT4 enrichment and transient β-catenin activation in stem cells (8 h–3 d), followed by signal decline by day 6.

### FLASH-RT Attenuates the Pro-Inflammatory CD4⁺ T-Cell Response during Intestinal Recovery

Given that FLASH promotes intestinal stem cell regeneration through Mrc1⁺ macrophage-derived WNT4 and creates a pro-regenerative innate immune microenvironment, we next asked whether the adaptive immune compartment likewise contributes to FLASH-induced intestinal recovery. To test this, we subclustered T cells from the scRNA-seq dataset, resolving three CD8⁺ T cell and three CD4⁺ T cell subsets (Fig. 5A, 5B). Among CD4⁺ T cells, we recognized a pro-inflammatory subset (CD4TC2) expressed Th1 markers Tnf and Tbx21 ^37^ and was less abundant in FLASH-irradiated groups than CONV (Fig. 5C, 5D). A second CD4^+^ subset (CD4TC1) expressed stem-cell-like/naive markers Tcf7 and Lef1 and was also reduced in FLASH groups (Fig. 5C, 5E). At day 3, C3 showed higher proportions of both root CD4TC1 and terminal CD4TC2 than F3. Pseudotime analysis placed CD4TC1 at the trajectory root and CD4TC2 at the terminal end, defining a unidirectional CD4TC1 → CD4TC2 differentiation path (Fig. 5F).

**Fig. 5.**
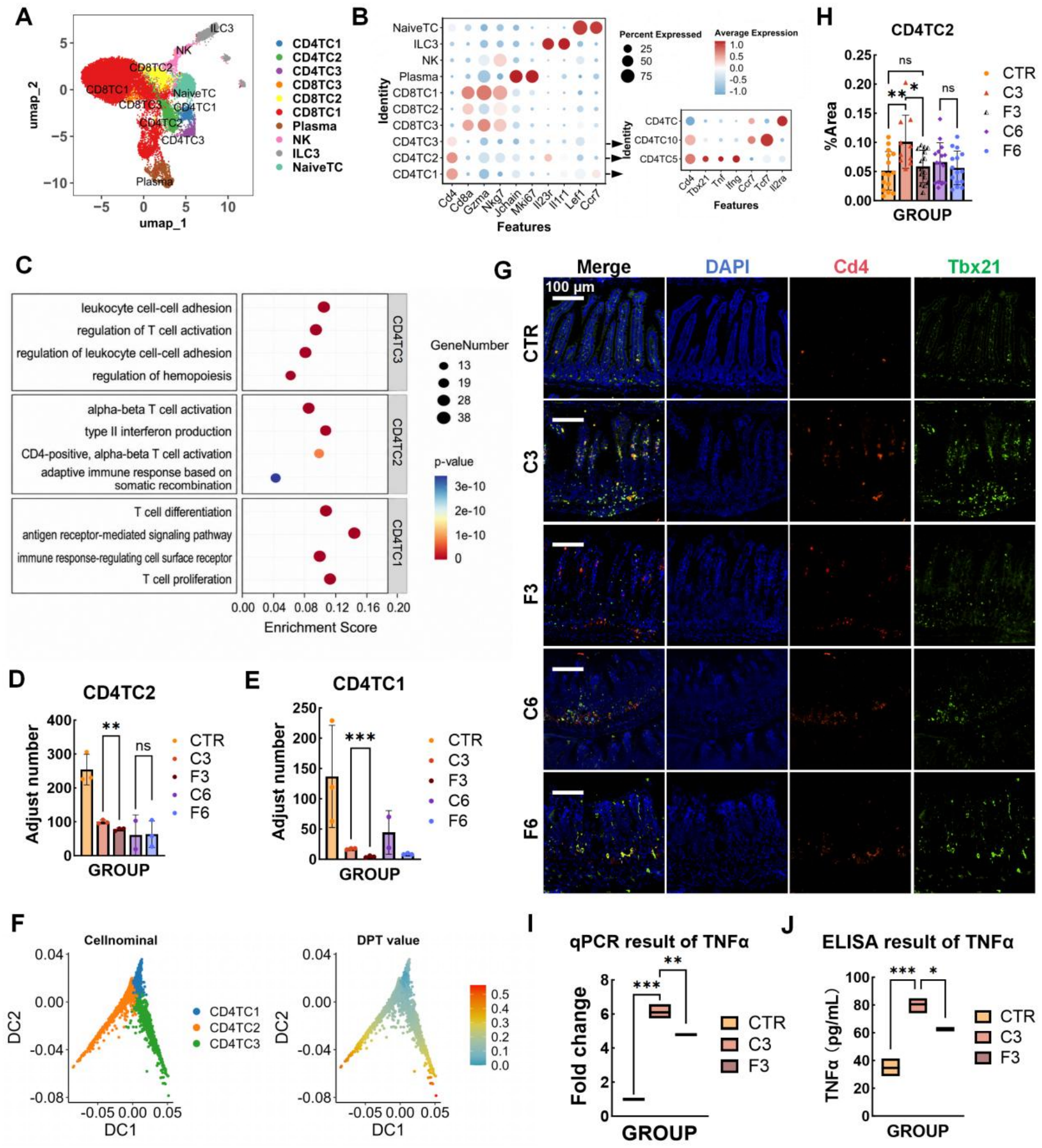
FLASH-RT remodels CD4⁺ T-cell states and attenuates inflammatory responses during intestinal regeneration. (A) UMAP visualization of intestinal T-cell populations identified by scRNA-seq. (B) Dot plot showing representative marker gene expression across the identified T-cell populations. (C) Gene Ontology (GO) enrichment analysis of differentially expressed genes in the indicated CD4⁺ T-cell populations. (D) Quantification of the CD4TC2 population across the indicated experimental groups (n=3). (E) Quantification of the CD4TC1 population across the indicated experimental groups (n=3). (F) Pseudotime trajectory analysis of intestinal CD4⁺ T-cell populations. DPT, diffusion pseudotime. (G) Representative multiplex immunofluorescence images showing CD4TC2 cells in the indicated experimental groups. (H) Quantification of CD4TC2 cells by multiplex immunofluorescence (n = 15 from three mice per group). (I) Relative Tnf mRNA expression in intestinal tissues from the indicated experimental groups, determined by quantitative PCR (qPCR) (n=3). (J) TNF-α levels in serum from the indicated experimental groups, determined by ELISA (n=3). Data are presented as mean ± SEM. Statistical significance was assessed by a one-way ANOVA followed by Tukey’s multiple comparisons test. *\*P* < 0.05, *\*\*P* < 0.01, *\*\*\*P* < 0.001. Scale bars are indicated.

To validate the above single-cell findings *in situ*, we quantified the infiltration abundance of Tbx21⁺ CD4⁺ cells in the intestinal mucosa using mIHC. Statistical analysis (Fig. 5G, 5H) showed that CD4TC2 in intestinal mucosa showed elevated infiltration in C3 relative to CTR (P < 0.01), whereas F3 was reduced relative to C3 (P < 0.05) and comparable to CTR; no significant differences were observed at day 6. To determine whether these cellular differences were accompanied by changes in the inflammatory milieu,, qPCR at 24 h after irradiation showed lower mRNA expression of pro-inflammatory cytokine TNF-α in FLASH-irradiated intestines than in CONV (Fig. 5I), and ELISA at 72 h confirmed lower serum TNF-α in FLASH-irradiated mice (Fig. 5J).

Together, these findings show that FLASH-RT is associated with a less pronounced pro-inflammatory CD4⁺ T-cell response during the early phase after irradiation, characterized by reduced abundance of both Tcf7⁺ less-differentiated and Tbx21⁺ inflammatory CD4⁺ T-cell states and lower TNF-α levels. Together with the enrichment of Mrc1⁺ macrophages described above, these results reveal coordinated remodeling of innate and adaptive immune states following FLASH-RT.

## Discussion

FLASH-RT reduces normal-tissue toxicity while preserving tumor control, yet the biological basis for this dose-rate effect remains incompletely defined. Here, single-cell transcriptomics across injury and recovery stages revealed that FLASH-RT and CONV-RT elicit distinct epithelial and immune responses. FLASH-RT preserved Olfm4⁺ stem cells, enriched Mrc1⁺ macrophages, and reduced Tbx21⁺ CD4⁺ T cells with diminished TNF-α production. Cell–cell communication analysis further identified enhanced WNT4 signaling from Mrc1⁺ macrophages toward Olfm4⁺ epithelial cells accompanied by transient β-catenin activation in early regeneration (Fig. 4). These findings indicate that FLASH-mediated intestinal protection reflects coordinated remodeling of epithelial regeneration and the immune microenvironment.

Preservation of regenerative epithelial populations provides an important cellular basis for the enhanced tissue recovery after FLASH. Our scRNA-seq data showed greater representation of Olfm4⁺ regenerative epithelial cells after FLASH-RT, with transcriptional features of active epithelial regeneration (Fig. 2B, 2C). This aligns with recent proton FLASH studies ^23^ showing preservation of Lgr5⁺ CBCs and injury-responsive regenerative stem cell populations characterized by Ly6a, Clu, Areg, and Anxa2 expression. However, that study primarily addressed epithelial cell-intrinsic oxidative stress and cell cycle changes, rather than remodeling of the immune microenvironment. Our results extend this observation by showing that FLASH preserves regenerative epithelial populations that subsequently expand during tissue recovery (Fig. 2F, 2G). Thus, FLASH may not simply spare more epithelial cells from radiation, but preserve a pool of cells capable of reinitiating and sustaining regenerative programs after injury.

Epithelial preservation was accompanied by substantial immune remodeling. Previous single-cell profiling of conventional radiation-induced intestinal injury revealed pronounced early myeloid remodeling, characterized by enrichment of Ly6c⁺ monocytes and pro-inflammatory macrophages around day 3, followed by recovery of resident macrophage populations ^30^. In contrast, a prominent myeloid feature associated with FLASH-RT in our study was the enrichment of Mrc1⁺ macrophages, which was further confirmed by mIHC (Fig. 3F, 3G). CellChat analysis predicted enhanced WNT4 communication from Mrc1⁺ macrophages toward Olfm4⁺ stem cell population (Fig. 3C), and mIHC supported spatial colocalization of WNT4 with these populations (Fig. 4F, 4G). These observations are consistent with previous work identifying macrophages as an important source of WNT signals during intestinal regeneration. Saha et al. ^38^ demonstrated that macrophage-derived extracellular vesicle-associated WNTs promote intestinal stem cell survival and recovery after radiation. Subsequent work demonstrated that macrophage β-catenin activity regulates WNT production, providing a feedback link between macrophage state and regenerative signaling ^39^. Thus, our findings place the FLASH-associated Mrc1⁺ macrophages enrichment within an established regenerative signaling framework, suggesting that radiation dose rate may influence not only epithelial survival but also the cellular source and availability of regenerative WNT signals.

The identity of the responding WNT receptor adds specificity to this model. WNT signaling depends not only on ligand abundance but also on receptor and coreceptor context. Ring et al. ^40^ systematically tested canonical and noncanonical WNT ligands to activate β-catenin-dependent transcription in different FZD receptor contexts. Notably, WNT4, despite its conventional classification as a noncanonical WNT ligand, was capable of inducing TCF/LEF-dependent transcription when paired with FZD5, FZD4, or FZD10, with LRP6 contributing as a required coreceptor. These findings establish a mechanistic basis for WNT4-dependent β-catenin signaling in an appropriate receptor context. In light of this work, our observation that FLASH enriches Mrc1⁺ macrophages while increasing predicted WNT4–FZD5 communication with Olfm4⁺ stem cells (Fig. 4A, 4G) provide a biologically plausible link between macrophage remodeling and stem cell signaling after FLASH. Importantly, CellChat identifies a predicted ligand–receptor interaction and the spatial colocalization supports cellular association. Together with prior mechanistic evidence for WNT4–FZD5-dependent β-catenin signaling and our downstream measurements, these data support a model in which Mrc1⁺ macrophages engage WNT4–β-catenin signaling during FLASH-associated intestinal regeneration.

The temporal behavior of this pathway adds a further dimension. WNT4 signaling became apparent as early as 8 h after FLASH, with β-catenin activation sustained through 24 h and day 3 but absent by day 6 (Fig. 4I). This transient pattern coincided with changes in WNT/β-catenin-responsive and stem cell-associated genes, including up-regulation of Axin2, Myc, and Tcf7 and down-regulation of Sfrp1 (Fig. 4J). These findings are consistent with an early role for WNT4-associated signaling in maintaining or reactivating regenerative stem cell programs following radiation. Rather than continuously elevating WNT signaling throughout repair, FLASH appears to activate the pathway within a defined regenerative window. This interpretation is reinforced by evidence that WNT4 has stage-specific functions during intestinal regeneration. A recent study ^36^ reported that, at later stages of radiation-induced intestinal regeneration, WNT4 suppressed canonical WNT/β-catenin signaling in organoids while promoting crypt symmetric fission through ROR2. Thus, WNT4 may function not as a single linear pathway but as a context-dependent signal whose output changes with receptor availability and tissue state. Our observation of early, transient β-catenin activation after FLASH is compatible with a model in which WNT4-FZD5 signaling contributes to the initial recovery of regenerative stem cell populations, whereas later WNT4–ROR2 signaling may participate in crypt architectural remodeling. Although our study does not establish a temporal receptor switch from FZD5 to ROR2, the distinct timing and reported functions of these modes provide a testable framework. Future experiments combining time-resolved analysis with receptor-specific perturbation will be important for determining whether FZD5- and ROR2-dependent WNT4 signaling make temporally distinct contributions to regeneration.

FLASH-associated immune remodeling extended beyond macrophages to the adaptive immune compartment. We observed reductions in both Tcf7⁺ and Tbx21⁺ CD4⁺ T-cell populations after FLASH, accompanied by decreased TNF-α production (Fig. 5I, 5J). T helper cell-derived cytokines regulate intestinal stem cell renewal and differentiation ^41^, while excessive CD4⁺ T-cell inflammation has been implicated in radiation induced intestinal injury. In particular, total-body irradiation increases intestinal CD4⁺ T-cell abundance and TNF-α expression, with concomitant damage to crypt stem cells and Paneth cells ^42^. In this context, reduce Tbx21⁺ CD4⁺ T cells and TNF-α production observed after FLASH (Fig. 5I, 5J) may reflect a less inflammatory immune environment during epithelial recovery. The pseudotemporal relationship between Tcf7⁺ and Tbx21⁺ CD4⁺ T-cell states further suggests that radiation dose rate may be associated with dynamic changes in CD4⁺ T-cell states during tissue repair, although this inferred trajectory does not establish lineage conversion. Thus, our findings identify an attenuated pro-inflammatory CD4⁺ T-cell response as a feature of the immune remodeling associated with FLASH irradiation during intestinal recovery..

Taken together, these findings support a model in which FLASH-associated intestinal recovery coincides with coordinated remodeling of regenerative and inflammatory immune states. On the myeloid side, FLASH-RT was associated with enrichment of Mrc1⁺ macrophages and increased WNT4-associated signaling involving Olfm4⁺ stem cells, accompanied by early and transient β-catenin activation during epithelial recovery (Fig. 2-4). On the lymphoid side, following FLASH-RT, Tbx21⁺ CD4⁺ T-cell abundance and TNF-α levels were reduced, consistent with an attenuated pro-inflammatory CD4⁺ T-cell response during intestinal recovery. (Fig. 5). These changes occurred within a similar early recovery window, suggesting that epithelial regeneration after FLASH-RT cannot be separated from the surrounding immune environment. Importantly, this framework does not replace the established view that FLASH protects normal tissue by reducing radiation-induced injury. Rather, it suggests that radiation dose rate may also influence the cellular and signaling landscape during subsequent recovery, including changes in immune-cell states and the regenerative signals accompanying epithelial repair.

From this perspective, the biological consequences of dose rate may depend not only on the magnitude of the initial radiation injury but also on the trajectory through which damaged tissue enters and sustains a regenerative state. The enrichment of Mrc1⁺ macrophages, together with WNT4-associated signaling involving Olfm4⁺ stem cells and transient β-catenin activation, provides a framework linking dose rate to regenerative niche remodeling, whereas the concurrent reduction in T-cell-associated inflammatory signaling reflects a distinct dimension of immune remodeling during recovery. Together, these observations position dose rate not merely as a physical determinant of radiation injury but also as a biological regulator of post-irradiation immune–stem cell interactions and tissue repair trajectories. More broadly, defining how radiation parameters shape regenerative cellular networks may provide a complementary framework for understanding normal-tissue sparing and for identifying strategies that enhance tissue recovery after radiotherapy.

Several aspects of this work warrant further investigation. First, while our transcriptomic analyses strongly implicate macrophage-derived signaling in stem-cell regeneration, the direct causality and downstream receptor dynamics of this immune–epithelial crosstalk require further functional validation. Second, the precise mechanisms by which FLASH-RT modulates CD4⁺ T-cell states, and the extent to which these specific immune alterations contribute to overall tissue recovery, remain to be fully elucidated.

## Materials and Methods

### Animal irradiation experiments

This study was designed to evaluate the protective effects of FLASH-RT on normal intestinal tissue. Female C57BL/6J mice (6–8 weeks old; strain code N000013) were purchased from GemPharmatech Co., Ltd. (Nanjing, China). Animals were housed in standardized cages under a 12-h light/12-h dark cycle with ad libitum access to standard chow and acidified drinking water. All animal experimental procedures were approved by XXX (Approval No. XXX) and were performed in accordance with the relevant guidelines and regulations.

For irradiation, mice were anesthetized and immobilized before exposure to a vertically oriented electron beam generated by a 6-MeV electron linear accelerator. For whole-abdomen irradiation (WAI), a collimator was installed at the accelerator exit window to define a 45 × 45 mm irradiation field. Lead shielding was used to protect the thorax, head and neck, and limbs from irradiation. The CONV-RT group received irradiation at a dose rate of 0.05 Gy/s, whereas the FLASH-RT group was irradiated at 750 Gy/s. Both groups received a total dose of 15 Gy. Sham-irradiated control mice underwent the same anesthesia and immobilization procedures but were not exposed to radiation.

### Single-cell suspension preparation and library construction

Fresh mouse small intestine tissues were collected under sterile conditions, immediately placed in prechilled phosphate-buffered saline (PBS), and rapidly transported to the laboratory for subsequent processing. Tissue samples were enzymatically dissociated with collagenase IV and DNase I at 37 °C for 30 min, with gentle pipetting during digestion to facilitate tissue dissociation. The resulting cell suspensions were sequentially passed through 40-μm and 70-μm cell strainers to remove undigested tissue debris. Cell viability was assessed by trypan blue exclusion, and samples with a viability of >85% were used for subsequent processing. Dead cells were removed using a Dead Cell Removal Kit. The resulting single-cell suspensions were adjusted to a concentration of approximately 700–1,200 cells/μL and processed using the 10x Genomics Chromium Controller with the Single Cell 3′ v3.1 Reagent Kit for single-cell partitioning, reverse transcription, and cDNA amplification. The resulting sequencing libraries were quantified using a Qubit 3.0 Fluorometer, and fragment size distributions were assessed using an Agilent 2100 Bioanalyzer. Libraries were subsequently sequenced on an Illumina NovaSeq 6000 platform using paired-end 150-bp (PE150) sequencing, with a target sequencing depth of approximately 50,000 reads per cell.

### Data preprocessing and quality control

Raw sequencing data in FASTQ format were processed using Cell Ranger (version 6.1.2) for sample demultiplexing, read alignment, and gene-level quantification. Reads were aligned to the mouse mm10 reference genome. The resulting gene expression matrices were imported into R and subjected to downstream analysis using Seurat (version 4.3.0). Cells were retained if they contained >200 and <7,500 detected genes (nFeature_RNA), >500 and <25,000 total UMI counts (nCount_RNA), and <25% mitochondrial gene-derived transcripts (percent.mt). Potential doublets were identified and removed using *DoubletFinder*. Following quality control, the resulting high-confidence single-cell transcriptomic profiles were used for downstream analyses.

### Dimensionality reduction, clustering, and differential expression analysis

Gene expression matrices were normalized using the “LogNormalize” method with a scale factor of 10,000. The 2,000 most highly variable genes were identified using the *FindVariableFeatures* function and subsequently used for principal component analysis (PCA). Uniform Manifold Approximation and Projection (UMAP) was performed for nonlinear dimensionality reduction and visualization. Cell–cell neighborhood graphs were constructed based on the first 40 principal components using *FindNeighbors*, followed by unsupervised clustering with *FindClusters* at a resolution of 0.9. Differentially expressed genes were identified using the *FindAllMarkers* or *FindMarkers* functions with the Wilcoxon rank-sum test. Genes with an absolute log2 fold change (|log2FC|) > 0.25 and an adjusted P value < 0.05 were considered differentially expressed. Cell identities were assigned based on established marker-gene expression profiles, reference annotations from the CellMarker database, and published literature. Gene Ontology (GO) and Kyoto Encyclopedia of Genes and Genomes (KEGG) enrichment analyses were performed using the clusterProfiler R package, with adjusted P < 0.05 considered statistically significant.

### CellChat analysis

Cell–cell communication within the mouse small intestinal microenvironment was investigated using CellChat (version 2) in R. Analyses were performed on quality-controlled and cell-type-annotated populations, with the average gene expression profile of each cell population used as input. The CellChatDB.mouse database was used as the reference ligand–receptor interaction database, incorporating experimentally and computationally supported ligand–receptor pairs, including multimeric ligand–receptor complexes. Overexpressed ligands and receptors were first identified, followed by estimation of intercellular communication probabilities using the *computeCommunProb* function. Communication probabilities for individual ligand–receptor pairs were inferred using a truncated-mean (*truncatedMean*) approach, and statistical significance was assessed by permutation testing with 100 permutations.

Ligand–receptor interactions were subsequently aggregated at the signaling-pathway level using the *computeCommunProbPathway* function to identify potentially relevant signaling pathways. The overall communication network was constructed using *aggregateNet*, and the relative roles of individual cell populations as signal senders (outgoing) and receivers (incoming) were evaluated using *netAnalysis_signalingRole_scatter* and *netAnalysis_signalingRole_heatmap*. Key intercellular signaling networks, including WNT4-mediated communication, were visualized using *netVisual_aggregate* and *netVisual_heatmap*. Unless otherwise specified, all statistically significant ligand–receptor interactions were considered, and interactions with a communication probability >0.05 were retained for downstream visualization and analysis.

### Pseudotime analysis and trajectory inference (Monocle 2 and DiffusionMap)

To characterize developmental dynamics and differentiation trajectories within selected cell lineages, pseudotime analysis was performed by integrating the reverse graph embedding framework implemented in Monocle 2 with the diffusion-map approach.

### Monocle 2 trajectory reconstruction and pseudotime ordering

Following quality control and cell-type annotation, the expression matrix of the lineage of interest was extracted for trajectory analysis. A *CellDataSet* object was constructed using the Monocle R package (version 2.26.0), followed by normalization and dispersion estimation using the *estimateSizeFactors* and *estimateDispersions* functions, respectively. To reduce dimensionality and reconstruct the underlying developmental trajectory, the DDRTree (Discriminative Dimensionality Reduction via Trees) algorithm was implemented using the *reduceDimension* function, with *max_components* set to 2 to define the principal graph structure.

Cells were ordered along the inferred trajectory using the *orderCells* function. The root state of the trajectory was manually assigned based on established biological markers and cellular functional states, with cell populations exhibiting relatively undifferentiated or progenitor-like characteristics (e.g., high expression of selected proliferation-associated markers) designated as the trajectory origin (*root_state*). Each cell was subsequently assigned a relative pseudotime value representing its progression from the inferred initial state toward more differentiated terminal states.

### DiffusionMap-based validation of global trajectory structure

To provide an orthogonal and less model-dependent assessment of the global topology of the differentiation landscape and to evaluate the robustness of the Monocle 2-derived pseudotime ordering, diffusion-map analysis was performed using the *destiny* R package (version 3.14.0). Briefly, pairwise Euclidean distances between cells were transformed into a Gaussian kernel, and the kernel width (σ) was automatically estimated using the *find_dm_k* function. This approach was used to capture the distribution of cells within a continuous latent-state space. The first two to three diffusion components (DCs) were extracted for dimensionality reduction and visualization, allowing the continuity of cellular states to be assessed independently of the Monocle 2 framework. The cellular distribution and global topology identified by *DiffusionMap* were then compared with the DDRTree trajectories reconstructed by *Monocle 2* to assess the consistency and robustness of the inferred differentiation paths.

### Hematoxylin and eosin (H&E) staining

Paraffin-embedded tissue sections were routinely deparaffinized and rehydrated. Briefly, sections were immersed sequentially in xylene I and II for 15 min each, followed by graded ethanol solutions (100% ethanol I and II, 95%, 85%, and 75%) for 5 min each, and then rinsed with distilled water. Sections were stained with hematoxylin for 8 min and rinsed under running water to remove excess stain. Sections were subsequently differentiated briefly in 1% hydrochloric acid ethanol until the background became clear, followed by washing under running water to terminate differentiation. Nuclei were blued in warm water for 10 min and rinsed with running water. Sections were then counterstained with 0.5% water-soluble eosin for 3 min and briefly rinsed with distilled water. After dehydration through a graded ethanol series (75%, 85%, 95%, and 100% ethanol I and II), sections were cleared in xylene and mounted with neutral resin. Tissue morphology was examined by light microscopy, and images were acquired using an Axio Scan.Z1 automated digital slide scanning system (Zeiss; 17026728).

### Immunohistochemistry (IHC)

Paraffin-embedded tissue sections were prepared using standard histological procedures. Briefly, tissue specimens were fixed in 4% paraformaldehyde, routinely dehydrated and cleared, and embedded in paraffin. Serial sections were cut at a thickness of 4 μm. Sections were deparaffinized in xylene and rehydrated through a graded ethanol series. Heat-induced antigen retrieval was performed by microwave treatment in either sodium citrate buffer (pH 6.0) or Tris-EDTA buffer (pH 9.0), according to the requirements of the respective primary antibody. Endogenous peroxidase activity was blocked by incubation with 3% hydrogen peroxide for 10 min at room temperature in the dark. Sections were subsequently blocked with 5% bovine serum albumin (BSA) in PBS for 30 min at room temperature to reduce nonspecific binding. Sections were incubated with the appropriate diluted primary antibody overnight at 4 °C in a humidified chamber. On the following day, sections were equilibrated to room temperature for 1 h and extensively washed with PBS before incubation with horseradish peroxidase (HRP)-conjugated secondary antibody [Goat Anti-Rabbit IgG H&L (HRP), 511203] for 1 h at room temperature. Following PBS washes, immunoreactivity was visualized using 3,3′-diaminobenzidine (DAB) as the chromogenic substrate. Sections were counterstained with hematoxylin, dehydrated through graded ethanol, cleared in xylene, and mounted with neutral resin.

### Multiplex immunohistochemistry (mIHC)

Multiplex staining was performed using a tyramide signal amplification (TSA)-based multiplex immunofluorescence protocol, enabling sequential labeling of multiple targets on a single paraffin-embedded tissue section. Briefly, paraffin sections were baked at 60 °C for 1 h, followed by deparaffinization in xylene and rehydration through a graded ethanol series. Heat-induced antigen retrieval was performed by microwave treatment in sodium citrate buffer (pH 6.0) or Tris-EDTA buffer (pH 9.0), followed by cooling to room temperature. Endogenous peroxidase activity was quenched by incubation with 3% hydrogen peroxide for 10 min in the dark. Sections were subsequently blocked with 5% BSA for 30 min to minimize nonspecific binding. The pretreatment steps before primary antibody incubation were identical to those described for conventional IHC.

For multiplex staining, targets were labeled sequentially according to a predefined staining order, with one target processed per staining cycle. Briefly, sections were incubated with the corresponding primary antibody overnight at 4 °C in a humidified chamber. After extensive washing with PBST, sections were incubated with an HRP-conjugated secondary antibody for 50 min at room temperature, followed by incubation with the corresponding TSA fluorophore working solution for 15 min in the dark, allowing covalent deposition of the fluorescent signal around the target antigen. After completion of each staining cycle, sections underwent microwave treatment under the same antigen-retrieval conditions to remove noncovalently bound antibody complexes while preserving the covalently deposited TSA signal. This cycle was repeated sequentially for all target antigens. After completion of the final staining cycle, nuclei were counterstained with DAPI, and sections were mounted using an antifade mounting medium. Images were acquired using a high-resolution digital pathology slide scanning system (ZCKP020190900100108).

The primary antibodies used were anti-Ki67 (Abcam, ab16667), anti-Fabp1 (Abcam, ab171739), anti-CD68 (Abcam, ab283654), anti-WNT4 (Bioss, bs-6134R), anti-EpCAM (CST, 93790T), anti-CD4 (CST, 25229T), anti-CD206 (CST, 24595T), anti-Olfm4 (CST, 39141T), anti-Ki67 (Abcam, ab16667), and anti-TBX21 (CST, 97135T). The secondary antibody was Goat Anti-Rabbit IgG H&L (HRP) (511203). Multiplex staining was performed using the Goat Anti-Mouse/Rabbit Multiplex IHC Detection Kit (Zhengneng Bio, 18005).

### Irradiation and recombinant mouse WNT4 treatment

Following conventional irradiation, mice received daily intraperitoneal injections of 200 μL recombinant mouse WNT4 protein (rmWNT4; 200 ng per mouse; TargetMOI, TMPH-04317) for 6 consecutive days. Mice in the corresponding control groups received an equal volume of PBS containing 1% BSA. Mice were euthanized on day 6 after irradiation, and small intestinal tissues were collected for subsequent analyses.

### AAV treatment experiment

For AAV-mediated WNT4 overexpression, mice received intraperitoneal administration of AAV-Wnt4 (AAV-CMV-Wnt4-FLAG-GdGreen; OBIO Technology Corp., Ltd., Shanghai, China) or the corresponding empty-vector control (AAV-CMV-GdGreen; OBIO Technology Corp., Ltd., Shanghai, China) 2 weeks before irradiation. A dose of 2 × 10^11^ vector genomes (vg) per mouse was administered. Mice were subsequently subjected to 15-Gy conventional abdominal irradiation. At the indicated time points after irradiation, mice were euthanized, and small intestinal tissues were collected for subsequent analyses.

### Western blotting

Total protein was extracted from tissue samples and cultured cells using ice-cold RIPA lysis buffer supplemented with a protease/phosphatase inhibitor cocktail (CST, 5872). Tissue samples were homogenized on ice using a tissue homogenizer for 5 min at −10 °C, whereas cultured cells were lysed in RIPA buffer followed by sonication. Lysates were incubated on ice and subsequently centrifuged to collect the supernatants. Protein concentrations were determined using a bicinchoninic acid (BCA) assay, and equal amounts of protein were loaded for electrophoresis. Protein samples were separated by 10% SDS–polyacrylamide gel electrophoresis and transferred onto polyvinylidene difluoride (PVDF) membranes.

Membranes were blocked with TBST containing 5% nonfat dry milk and incubated overnight at 4 °C with the indicated primary antibodies at a dilution of 1:1,000. After extensive washing with TBST, membranes were incubated with HRP-conjugated secondary antibody (1:2,000) for 1 h at room temperature. Protein signals were detected using an enhanced chemiluminescence (ECL) substrate and acquired using a chemiluminescence imaging system. Band intensities were quantified using ImageJ.

The primary antibodies used were anti-WNT4 (Bioss, BS-6134R), anti-β-catenin (CST, R381559), anti-Non-phospho (Active) β-catenin (Ser45) (CST, 19807T), anti-β-actin (Beyotime, AF5003), anti-α-actinin (CST, 6487T) and anti-Lamin B1 (CST, 13435T). The secondary antibody was HRP-conjugated goat anti-rabbit IgG (Proteintech, SA00001-2).

### Quantitative real-time PCR (qPCR)

Total RNA was extracted from intestinal tissue samples and reverse-transcribed into complementary DNA (cDNA). Quantitative PCR was performed using 2× Real Fast SYBR qPCR Premix. Relative mRNA expression levels were calculated using the comparative threshold cycle (2^−ΔΔCt^) method.

### Enzyme-linked immunosorbent assay (ELISA)

Serum samples were collected from three mice in each experimental group (CTR, CONV, and FLASH). Whole blood was allowed to clot at room temperature for 30 min and subsequently centrifuged at 6,000 rpm for 10 min. The resulting serum was collected and stored at −80 °C until analysis, with repeated freeze–thaw cycles avoided.

Serum TNF-α concentrations were measured using a high-sensitivity mouse TNF-α ELISA kit (MULTI SCIENCE, EK282) based on a sandwich ELISA format. Briefly, the kit reagents and serum samples were equilibrated to room temperature (25 °C). Standard, blank, and sample wells were prepared, and 100 μL of the corresponding standards or diluted samples was added to each well. Plates were covered and incubated at room temperature according to the manufacturer’s instructions. Following removal of the liquid and washing, biotinylated detection antibody working solution was added to each well and incubated at room temperature. After washing, HRP-conjugated avidin working solution was added and incubated at room temperature. Following another washing step, TMB substrate was added and incubated in the dark at room temperature. The reaction was terminated by addition of stop solution, and optical density (OD) was immediately measured at 450 nm using a microplate reader (BioTek Cytation 5). TNF-α concentrations were calculated from a standard curve generated from the relationship between standard concentrations and OD450 values. All samples were analyzed in technical duplicate, and the mean value was used for subsequent analyses.

## Statistical analysis

Data from independent validation experiments are presented as the arithmetic mean ± SEM. Statistical analyses and graphical representations were performed using GraphPad Prism 10.1.2 (San Diego, CA, USA). Data distributions were assessed using the Shapiro–Wilk test. For normally distributed data, comparisons between two groups were performed using an unpaired Student’s t test, whereas comparisons among multiple groups were performed using one-way analysis of variance (ANOVA) followed by Tukey’s post hoc multiple-comparison test. For data that did not satisfy the assumptions of normality, appropriate nonparametric tests were applied. A two-sided P value <0.05 was considered statistically significant. Statistical significance was denoted as follows: *P < 0.05, **P < 0.01, ***P < 0.001.

## Data, Materials, and Software Availability

Single-cell RNA sequencing data will be deposited at the NCBI’s Gene Expression Omnibus (GEO) database and made publicly available at the time of publication. The accession number is pending. This paper does not report original code. All other data are included in the article and/or SI Appendix.

## Acknowledgments

This work was supported by the National Natural Science Foundation of China (Grant No. 12441518) and the National Key Research and Development Program of China (Grant No. 2022YFC2402300).

## Author Contributions

H.L.and Q.F. designed research; Y.Z., Y.W., Y.Z., C.L., Y.W., C.H., C.L., J.W., H.Z., R.Q., Z.L. ,T.H and G.D. performed research; Y.Z., Y.Z. and C.L performed all irradiation and dissemination experiments; Y.Z., C.L. performed all immunofluorescence staining; Y.Z.,Y.Z. and C.L. performed H&E staining and data analysis; Y.W. and Y.Z. performed all RT-qPCR experiments and data analysis. Y.Z., Y.W. and C.L. performed single-cell RNAseq analyses; Y.Z., H.L. and Q.F. discussed data and commented on the manuscript; and Y.Z., H.L. and Q.F. wrote the paper, Y.Z., H.L., C.W., W.W. and Q.F. revised the paper.

